# Spatiotemporal control of gene expression boundaries using a feedforward loop

**DOI:** 10.1101/806620

**Authors:** Prasad U. Bandodkar, Hadel Al Asafen, Gregory T. Reeves

**Affiliations:** Department of Chemical and Biomolecular Engineering, North Carolina State University, Raleigh, North Carolina, United States of America; Department of Chemical Engineering and Applied Chemistry, University of Toronto, Toronto, CA; Program in Genetics, North Carolina State University, Raleigh, North Carolina, United States of America

**Keywords:** Feed Forward Loop, Dorsal morphogen, Twist, Spatiotemporal dynamics, Zelda, transcription factor, Drosophila embryo, Gene expression model

## Abstract

A feed forward loop (FFL) is commonly observed in several biological networks. The FFL network motif has been mostly been studied with respect to variation of the input signal in time, with only a few studies of FFL activity in a spatially distributed system such as morphogen-mediated tissue patterning. However, most morphogen gradients also evolve in time. We studied the spatiotemporal behavior of a coherent FFL in two contexts: (1) a generic, oscillating morphogen gradient and (2) the dorsal-ventral patterning of the early *Drosophila* embryo by a gradient of the NF-κB homolog Dorsal with its early target Twist. In both models, we found features in the dynamics of the intermediate node – phase difference and noise filtering – that were largely independent of the parameterization of the models, and thus were functions of the structure of the FFL itself. In the Dorsal gradient model, we also found that the dynamics of Dorsal require maternal pioneering factor Zelda for proper target gene expression.

## Introduction

In animal development, establishment of proper gene expression patterns within tissues is crucial to the fatemap and fitness of the organism. As such, regulation of gene expression is in place to ensure patterns are robust to perturbations in developmental conditions. While development is initiated by maternal morphogen gradients, robust gene expression often arises from the coordinated effort of several downstream transcription factors that are wired together to form gene regulatory networks. These networks may be represented as graphs, with genes represented by nodes and directed edges (arrows) representing direction of regulatory interactions. In such a network, there are some patterns, called “network motifs,” that seem to occur more frequently than others when compared with randomized networks.

In a motif described as a feed forward loop (FFL), transcription factor A regulates expression of two targets, B and C (Fig. 1A). B also acts as a transcription factor for C and transcriptional control is shared by both A and B. An FFL is called coherent (cFFL) when the direct effect of A on C and the indirect effect of A on C via B has the same sign, while it is called incoherent (iFFL) when the signs are opposite. These FFLs have been subjected to extensive mathematical analysis, both generally and in the context of real biological networks (Mangan and Alon, 2003; Mangan et al., 2003). However, these studies have been primarily restricted to studying the dynamics of FFLs in time upon transient or sustained changes to the input A. Since the FFL network motif occurs frequently and is found in several biological networks, it was expected that the structure of the motif serve a specific purpose. Indeed, cFFLs were found to delay the response of target genes to an increasing input (Mangan et al., 2003). This implies that detectable response of targets is feasible only when there is a sustained input signal and not when the signal is in short pulses. Such a mechanism would be effective in buffering small-time scale variations in the input signal.

**Figure 1.**
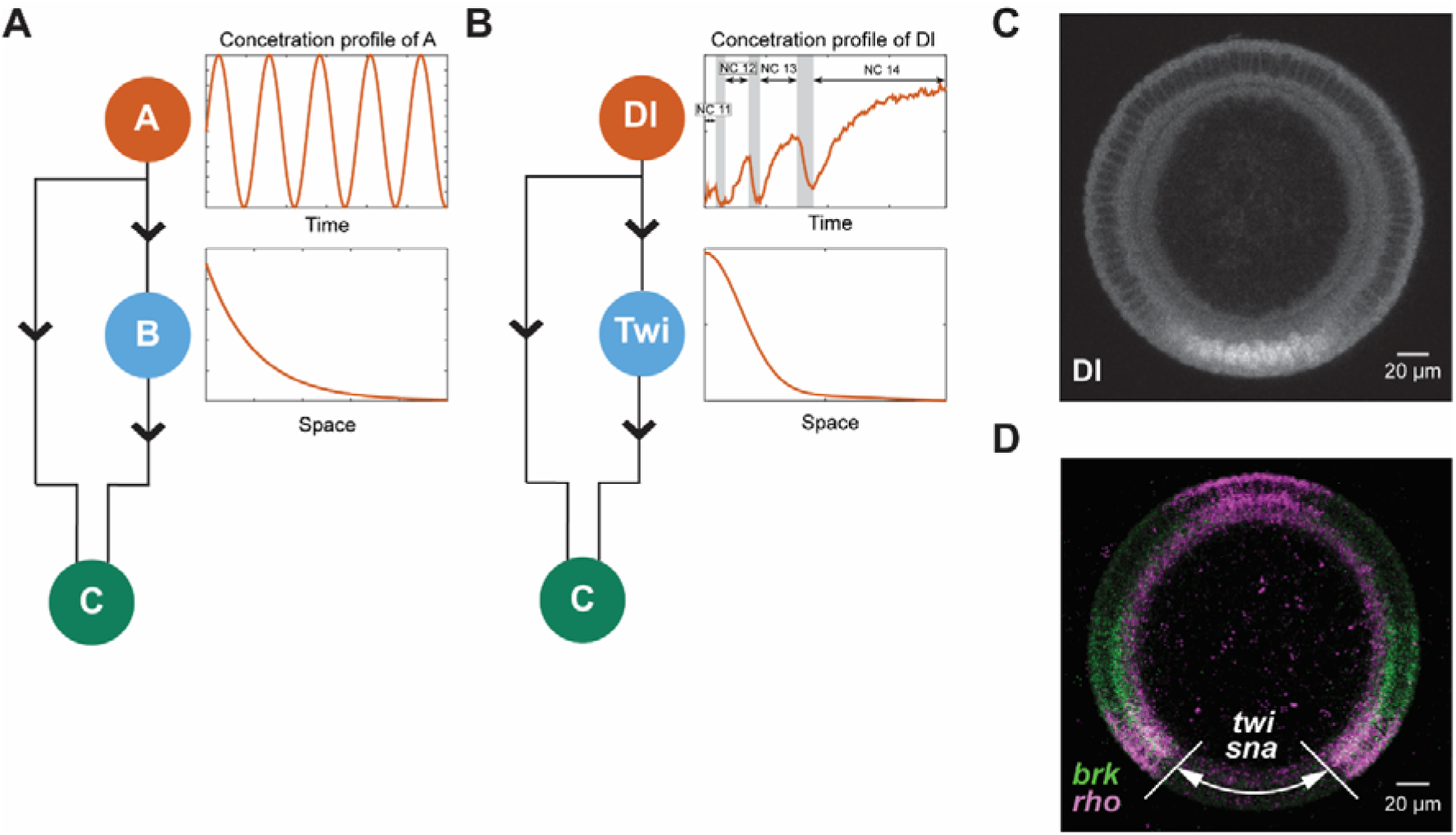
Introduction to the models. **A**. Schematic figure depicting the generic morphogen gradient model. The inset shows the variation of the concentration of the primary transcription factor A in space and time. **B**. Schematic figure depicting the Dl/Twi/Zld model. The inset shows the variation of the concentration of the primary transcription factor Dl in space and time. The peaks in the Dl gradient amplitude in time correspond to nuclear cycles 11-14. **C.** Fluorescence image of an anti-Dl immunostaining in a cross-sectioned embryo at nuclear cycle 14. **D.** Fluorescence image showing *brk* and *rho*, two genes with expression borders between 20% - 50% DV length. Domain of *twi* and *sna* expression annotated in white. Expression of *rho* on the dorsal 10% of the embryo is not Dorsal-dependent.

A smaller set of studies have analyzed the role of FFLs activated by morphogen gradients at steady state in tissue patterning (Zinzen et al., 2006). However, the field of developmental biology is discovering that morphogen gradients are rarely at steady state (DeLotto et al., 2007; Gregor et al., 2007; Heemskerk et al., 2019; Nahmad and Stathopoulos, 2009; Reeves et al., 2012; Ribes et al., 2010). Also, Sanger and Briscoe suggest that the level and duration of a morphogen signal, that might differ at various tissue locations, in combination with a network motif like an FFL can produce stable gene expression boundaries (Sagner and Briscoe, 2017). As such, some studies have also focused on the role of FFLs when the morphogen was varying both in space and time. Schaerli et al studied networks from the standpoint of stripe formation. They connected networks with 3 nodes with all possible combinations of regulation between them to predict stripe formation, and differentiated possible stripe-forming network topologies on the basis of similar dynamical mechanisms (Schaerli et al., 2014). They found that the minimal 3-node network to form stripes requires the motif to be incoherent in nature. In this study we investigated the function of coherent FFLs in both space and time to determine whether the cFFL could buffer gene expression against a time-varying morphogen gradient.

We first analyzed a general, dynamic morphogen gradient, which we modeled as having sinusoidal oscillations in time and an exponential decay in space (Fig. 1A). The generic morphogen, A, activates a secondary transcription factor, B. The two transcription factors then jointly activate the final target, C. We investigated whether the FFL may be more robust than if regulation were controlled by the initial activator (A) independently. We found that, under general conditions, the FFL indeed resulted in more stable (less oscillatory) gene expression boundaries than if A alone were activating C. The results were observed under a wide variety of parameter sets, indicating that the stabilization was a result of the structure of an FFL, rather than specific values of parameters. In particular, the intermediate transcription factor, B, had a modest phase difference from the oscillations in A, and furthermore, B generally acted as a noise filter on the dynamics of A. Both attributes of B contributed to the stabilization of C. After analysis of the generic morphogen model, we focused our analysis on the particular example of patterning of the dorsal-ventral (DV) axis of the early *Drosophila melanogaster* embryo by the morphogen Dorsal (Dl).

The Dl gradient has a bell-shaped distribution in space and is highly dynamic, with oscillations that are underscored by the relatively rapid syncytial nuclear divisions (Fig. 1B,C; (DeLotto et al., 2007; Liberman et al., 2009; Reeves et al., 2012)). As has been shown previously, the nuclear concentration of Dl varies significantly between nc 11 – nc 14 (Reeves et al., 2012), which, theoretically, may be incorrectly interpreted by the cells. It is possible that a cFFL through *twist* (*twi*), which is an early, high threshold target of Dl and codes for a transcription factor that activates gene expression with Dl, could potentially buffer fluctuations to result in the observed stability and consistency of target gene expression borders.

Several studies have indicated that genes expressed along the DV axis from the prospective mesoderm to the prospective ectoderm may be jointly regulated by the coordinated effort of Dl and Twi (Ip et al., 1992b, 1992a; Szymanski and Levine, 1995). Additionally, binding sites of Dl and Twi have been found in close proximity in the regulatory regions of these genes, suggesting that there could be synergistic interactions between the two proteins (Levine and Papatsenko, 2002; Ozdemir et al., 2011). Expression of these Dl/Twi target genes is limited to the ventral half of the embryo where the Dl and Twi gradients are active. Type I genes, such as *snail* (*sna*) and *twi* itself, are expressed on the ventral 20% of the embryo (Fig. 1D). Type II genes, such as *rhomboid* (*rho*), are repressed by Sna activity, and have dorsal borders around 35% DV (Fig. 1D). Finally, Type III genes, such as *brinker* (*brk*), are also repressed by Sna, but have domains that extend to roughly 50% DV (Fig. 1D).

We used Dl expression data (Reeves et al., 2012) as inputs to a model of the Dl/Twi cFFL. We modeled the expression of a target gene, C (Fig. 1B), which can have an expression border anywhere from 20% DV to 50% DV, as true Dl/Twi target genes do. As with the generic morphogen model, we investigated the stability of the gene expression border of the final target in the presence and absence of an FFL. We first found that, as a result of the short duration of nc 11-13, early activation of target genes was not possible. This problem also affected *twi*, which ruled out its role in a presumptive FFL with Dl. However, it is known experimentally that zygotic gene expression begins in nc 10 and observable gene expression for *twi* is first observed in nc 12, localized to the 20% ventral-most nuclei (Alberga et al., 1991; Jiang et al., 1991; Sandler and Stathopoulos, 2016; Thisse et al., 1988). Therefore, we included in our model the activity of pioneer factor Zelda (Zld), which has been shown to make enhancer regions accessible for transcription factors and to locally increase transcription factor concentration at enhancer sites (Dufourt et al., 2018; Liang et al., 2008; Mir et al., 2017; Nien et al., 2011; Yamada et al., 2019). A previous model of Zld activity showed that cooperativity between Zld and Dl may help explain Dl-dependent gene expression patterns (Kanodia et al., 2012).

After properly accounting for Zld activity, we found that, in the Dl/Twi system, in spite of the irregular Dl gradient amplitudes and oscillation periods, the case where Dl and Twi act together in an FFL outperforms the singular action of Dl. The observed stabilization in the boundary of target genes was due to a phase difference in the Dl and Twi concentrations that were largely independent of the parameter values characterizing the model. We also examined a *twi* mutant case, in which the target gene, C, is designed to respond to both Dl and Twi, yet Twi was absent. While the FFL often outperformed the *twi* mutant, shifting gene expression ventrally in the *twi* mutant helped to artificially stabilize gene expression. Together, the results of both the generic and the Dl/Twi models demonstrate that it is the structure of a coherent FFL that buffers oscillations in an input signal that varies both in space and time under variety of input conditions and parameter values.

## Results

### Generic morphogen gradient model

To study the effect of the cFFL on the dynamics of a morphogen system, we first investigated a model in which a generic morphogen, A, stably oscillates (sinusoidally) in time and has an exponential distribution in space (Fig. 1A, Fig. 2A,B; see Eqn 2 in Methods). The primary transcription factor A activates B, the intermediate node in the FFL, so that the gene expression border of B is located at roughly 20% tissue length (*x* = 0.2) on average (see Methods). Through a combination of diffusion and degradation, the secondary transcription factor B is also present in a concentration gradient, which varies slightly in time due to oscillations in A. The gradient of B is determined by its diffusive length scale (*λ*) and lifetime (*τ*; see Eqn 3 in Methods). Due to the oscillations in A, one may expect the concentration of B to also oscillate. However, the concentration profile of transcription factor B (simulated with parameter values *λ* = 0.1 and *τ* = 10) appears largely stable in time, except near the boundary of B expression (*x* = 0.2), where one might expect the profile of B to be the most sensitive (Fig. 2C). The oscillations in B near *x* = 0.2 are slightly out of phase with the oscillations in A (Fig. 2D), which may potentially impart stability to downstream gene expression.

**Figure 2.**
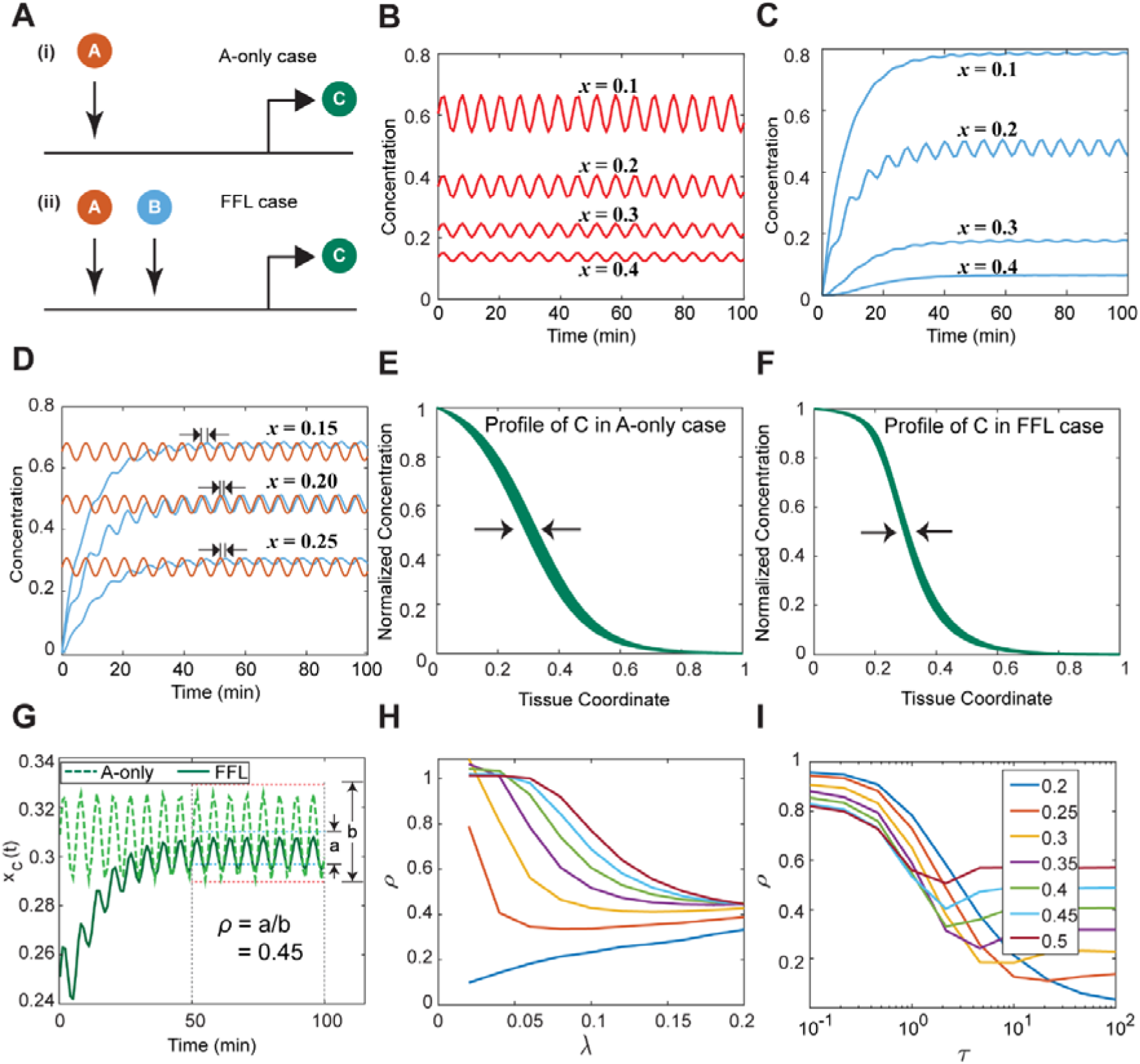
Generic morphogen gradient model. **A.** The enhancer module of C regulated by (i). A-only and by (ii) Feed Forward Loop (FFL). **B.** Variation of the concentration of primary transcription factor A in time, plotted at different values of *x*. **C.** Variation of the concentration of intermediate node B in time, plotted at different values of x. **D.** There is a slight phase difference between the concentration curves of A and B. This plot shows the variation of concentration of the intermediate node around its boundary (at *x* = 0.2) plotted against A, where the curves of A are representative of A at *x* = 0.2 that have been moved along the y axis to show the phase difference. **E.** Normalized concentration profile of C in the A-only case plotted after the initial transients have died. **F.** Normalized concentration profile of C in the FFL case plotted after the initial transients have died. Arrows in E and F highlight the extent of variation in space of the half-max of gene expression. **G.** Variation in the boundary of C against time for both cases. **H.** Variation of the extent of oscillations with at different values of *x_co_* **I.** Variation of the extent of oscillations with at different values of *x*_*c*0_.

We next analyzed the final target C, whose expression border in the absence of oscillations is located at *x* = *x*_*c*0_ (see Methods). We calculated the expression profile of C for two cases: (1) the “A-only” case, when C is designed to be activated only by A, and (2) the “FFL” case, when C is co-activated by A and B in an FFL. In both cases, the gene expression boundary of *C*, *x_C_* (*t*), oscillated in time due to the oscillations in A. However, by-eye it appeared that the expression boundary oscillated with a smaller amplitude in the FFL case than in the A-only case (compare Fig. 2E to F). To quantify whether the FFL stabilizes gene expression compared to the A-only case, we took the ratio of the extent of oscillations between the two cases (see Fig. 2G). If this ratio, which we call *ρ*, is less than 1, the FFL results in more stable boundaries. For the scenario shown in Fig. 2G, in which *λ* = 0.1, *ρ* = 10, and *x*_*c*0_ = 0.3, this ratio is = 0.45, which reflects the fact that the FFL visually results in oscillations with smaller amplitude. One possible explanation would be the phase difference in the concentrations of A and B, which can be seen clearly around the boundary of B and less clearly elsewhere (see Fig 2D and Movie S1-3).

To determine whether the stabilizing effect of the FFL was general, or perhaps limited to the particular choice of parameter values above, we calculated the metric for different genes with borders between *x*_*c*0_ = 0.2 and 0.5 while varying the parameters of the intermediate node B. We first varied (the diffusive length scale of B) and fixed the lifetime of B to be = 10. We found that the FFL stabilizes nearly all gene expression between *x*_*c*0_ = 0.2 and 0.5 (i.e., *ρ* <1 for those genes), irrespective of how far B diffuses (Fig. 2H). However, with the exception of the curve for *x*_*c*0_ = 0.2 which is very close to the boundary of B expression, the FFL tends to stabilize gene expression more when B is able to diffuse further. This implies that, a higher diffusive length scale of B results in better regulation of targets farther away. Thus, in the generic model, the FFL will almost always stabilize gene expression boundaries as long as the secondary transcription factor can diffuse to those boundaries. Since the tissue length is scaled to vary from 0 to 1, a value of = 0.1 is a reasonable choice for diffusive length scale of B, as it corresponds to a concentration gradient that extends roughly half-way around the tissue to include the domain of 0.2 < *x*_*c*0_ < 0.5 (Fig. S1).

Thus, holding *λ* = 0.1, we next varied the lifetime, *τ*, to determine its effect on the stabilizing role of the FFL. We found that for small values of *λ*, the FFL does little to improve on the independent action of transcription factor A (i.e., *ρ* ~ 1 Fig. 2I). As the value of *τ* increases, each gene curve of *ρ* as a function of *τ* drops below *ρ* = 1, and approaches a minimum before becoming flat. Since the oscillation period of A is 2π, the lifetime of the intermediate π node must be on roughly the same time scale or longer for the FFL to realize stable expression borders. If the lifetime of B is too short, the gradient of B simply oscillates with A, in phase. Once the lifetime of B is on the same scale as the oscillation period of A, B can act as a noise filter to stabilize gene expression boundaries. Much longer lifetimes also provide this benefit but may not be biologically realistic. This effect of is also illustrated visually in the normalized expression profiles of A, B and, C in the two models, namely A-only and FFL, plotted after the initial transients have decayed (see Movie S1 compared to Movie S3). We conclude that there is a reasonably wide range of diffusive length scales and protein lifetimes that result in stable gene expression boundaries, due in part to the phase differences between A and B and the noise-filtering of B (see also Fig. S2).

### The effect of Zelda on the Dorsal gradient and twist expression

After finding that, generally, the cFFL stabilizes gene expression boundaries in the face of a generic, oscillating morphogen, we then analyzed a model based upon experimentally obtained spatio-temporal data of Dl as the primary transcription factor (see Figs. 3A,B; (Reeves et al., 2012)). Unlike the generic morphogen, the oscillations of the Dl gradient do not have a regular period, even though they achieve limited oscillations due to syncytial nuclear divisions. During interphase, the amplitude of the Dl gradient constantly increases, then sharply decreases during mitosis. Moreover, the average amplitude within each interphase progressively increases from one nuclear cycle to the next, whether it be from each nuclear cycle being longer than the one prior, or some memory of previous state (Fig 3A,B). The highest levels of Dl are observed at the end of nc 14 interphase, which lasts about four times longer than the previous three nuclear cycle interphases (Fig 3B). Even so, the Dl gradient does not fully attain steady state by the end of nc 14. These precise dynamics are problematic for models of gene expression (O’Connell and Reeves, 2015; Reeves et al., 2012).

**Figure 3.**
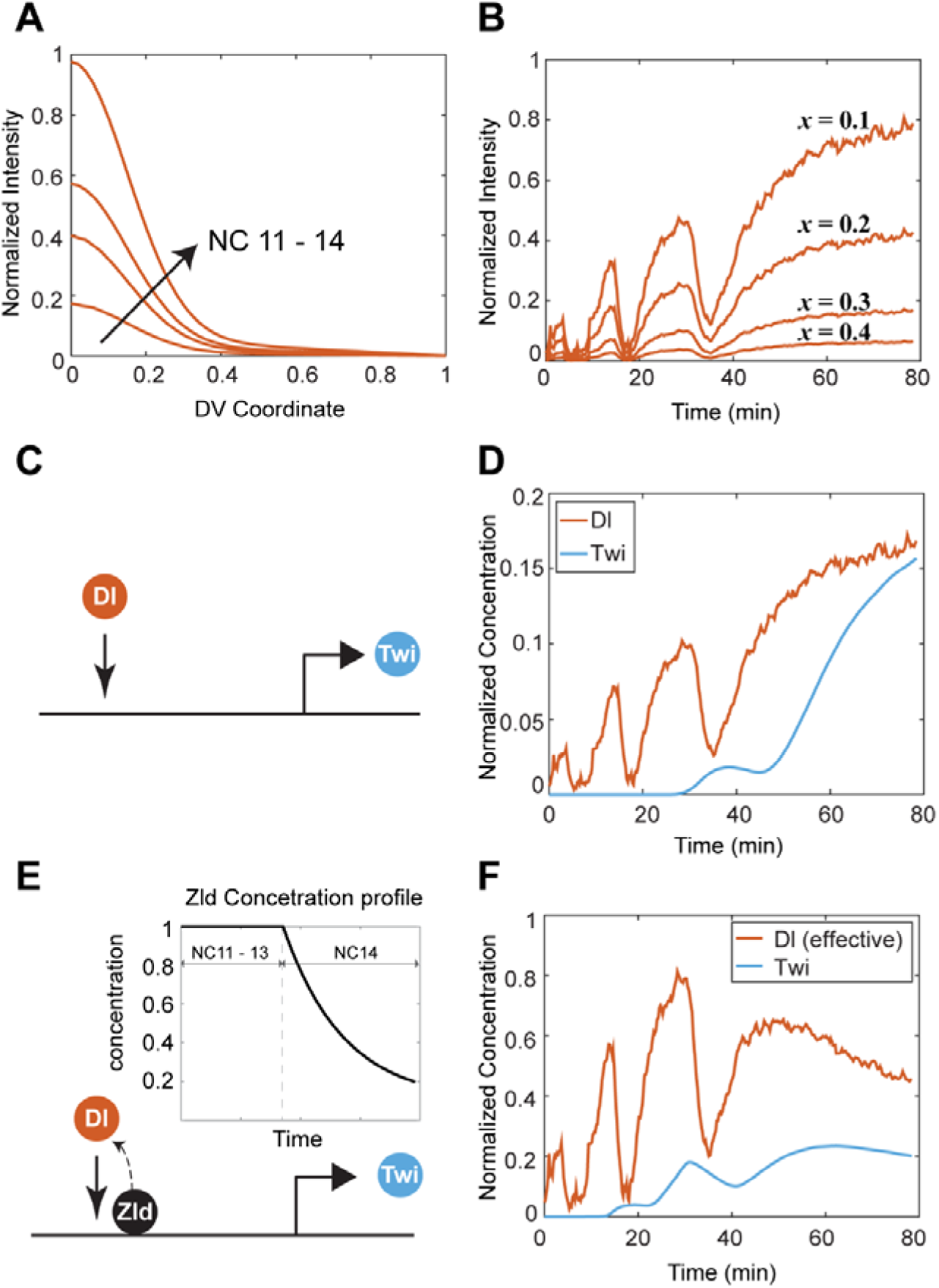
Effect of Zelda on the Dorsal gradient and Twist expression. **A.** Normalized Dl intensities as obtained from Venus-tagged Dl (Reeves et al., 2012) varying in space at the end of nuclear cycle 11 - 14. **B.** Normalized Dl intensities varying with time at different values of x. **C.** Schematic of the enhancer module of *twi* where Dl activates *twi.* **D.** Dl and Twi concentration curves at *x* = 0.3 for parameters = 0.1 and = 10. **E.** Schematic of the enhancer module of *twi* where Zld potentiates the activity of Dl and activates *twi.* Inset shows the dynamics of Zld concentration over time. **F.** Effective Dl concentration, as a result of Zld, and Twi concentration curves at *x* = 0.3 for parameters = 0.1 and = 10.

In the Dl gradient system, *twi* is activated by Dl (Fig. 3C) and together, Dl and Twi act in a cFFL (Ip et al., 1992b, 1992a; Szymanski and Levine, 1995). *twi* is expressed on the ventral 20% of the embryo (*x_Twi_* = 0.2), and *Twi* protein has been detected as early as late nc 12 (Alberga et al., 1991; Jiang et al., 1991; Sandler and Stathopoulos, 2016; Thisse et al., 1988). However, when we used the published Dl gradient data (Reeves et al., 2012) as the input to our gene expression model for *twi* (see Eqn 6 in Methods), the simulated *twi* activation was delayed until late nc 13 (Fig 3D). This delay resulted in insignificant *Twi* levels, compared to Dl levels, until mid-nc 14 at almost all relevant values of the DV coordinate (see Fig. S3). In our simulations, we found that the delayed activation of *Twi* rendered it practically ineffective as a collaborator in the FFL with Dl. In order to resolve the inconsistency in experimental observations and our gene expression model, we included the effect of the ubiquitous and pioneer maternal factor Zelda (Zld) (Liang et al., 2008; Nien et al., 2011; Sun et al., 2015).

Zld cooperatively interacts with Dl by enhancing its activity near the *cis*-regulatory modules (CRMs) of DV genes (Foo et al., 2014; Kanodia et al., 2012; Yamada et al., 2019). The cooperative effect of Zld on Dl has been previously modeled as an effective increase in the affinity of Dl for its DNA binding site (Kanodia et al., 2012). The increase in affinity could equivalently be seen as an increase in the effective Dl concentration by a factor *v*:[*DI*]*_effective_* = *v*[*DI*]. The factor is given by:

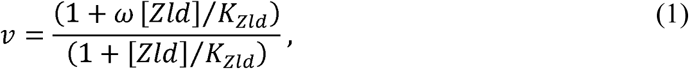

where *K_Zld_* refers to the DNA binding affinity of Zld and *ω* is a measure of Zld cooperativity with Dl. Thus, a high Zld binding affinity (low *K_Zld_*) or high cooperativity (high *ω*) leads to an increase in the effective Dl concentration, which leads to an increase in the activity of the regulatory region.

It is also important to note that Zld is a maternal protein that decays almost completely by the end of nc 14 (Fig. 3E), by which time transcription is entirely regulated by zygotic transcription factors. Dl levels on the other hand, have the steepest increase in concentrations in nc 14. It is expected that the effect of increasing levels of the graded factor Dl is offset by decreasing levels of the ubiquitous factor Zld, resulting in stable gene expression domains (Kanodia et al., 2012).

When we include cooperative interactions between Dl and Zld in the model (Fig. 3E), *twi* expression occurs earlier, at nc 12, which is consistent with experimental observations (Fig. 3F). Furthermore, the earlier expression of *twi* allows sufficient time for *Twi* levels to accumulate to enhance the performance of the FFL. An additional effect of Zld is to stabilize the Dl gradient: in the presence of Zld, the effective concentration of Dl in nc 13 becomes comparable to that in nc 14 (Fig. 3F; compare to Fig. 3D). There is a wide range of Zld model parameters (i.e, binding affinities and cooperativity) for which Dl activity in nc 13 and 14 are roughly similar (see Fig. S3).

### Downstream gene expression in the Dorsal/Twist/Zelda model

Given that *Twi* levels adequately represent experimental measurements when Zld is accounted for, we next simulated the dynamics of the output gene, C, which has a final expression border located at *x* = *x_c,f_* (see Methods). We chose the value of *x_c,f_* to lie between 0.2 and 0.5, where most Dl/Twi targets lie (Fig. 1D) and analyzed two cases similar to those in the generic model. In the first case (“Dl-only”; Fig. 4A(i)), the output gene, C, is hypothetically designed to only be activated by Dl. In the second case (“wildtype FFL”; Fig. 4A(ii)), C is regulated by both Dl and Twi, similar to the enhancer structure of many Dl target genes. Since Zld is a pioneer factor, it acts on both transcription factors Dl and Twi.

**Figure 4.**
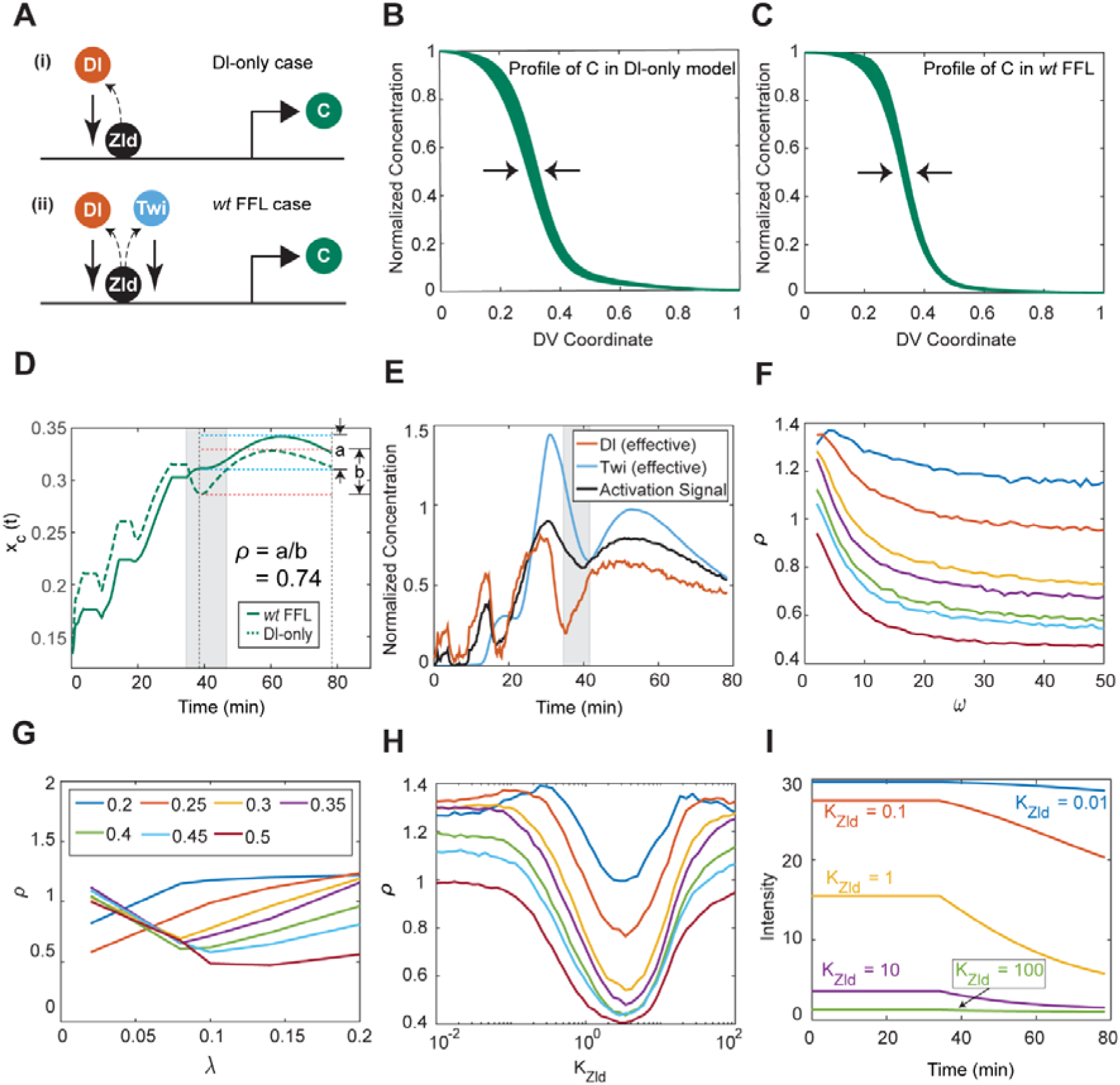
Effect of Dl-Twi FFL in the full model. **A.** The enhancer module of C regulated by (i). Dl-only and by (ii) Feed Forward Loop (FFL). Both cases include the effect of Zld. **B.** Normalized concentration profile of C in the Dl-only case plotted from ∼5 minutes into nc 14 till the beginning of gastrulation (∼80 min) **C.** Normalized concentration profile of C in the wildtype FFL case plotted from ∼5 minutes into nc 14 till the beginning of gastrulation (∼80 min). **D.** Variation in the boundary of C against time for both cases. **E.** Effective Dl and *Twi* concentration curves along with the Activation signal (see Eqn 9) at *x* = 0.3. **F.** Variation of the extent of oscillations with at different values of *x_c,f_* **G.** Variation of the extent of oscillations with at different values of *x_c,f_* **H.** Variation of the extent of oscillations with *K_Zld_* at different values of *x_c,f_* **I.** Variation of the effect of Zld (see Eqn. 1) in time at different values of *K_Zld_*.

In both cases (Dl-only and wildtype), the C expression profile varies in time in response to the varying concentrations of transcription factors Dl and *Twi* (Figs. 4B,C). Typically, the expression boundary of C – where the profile of C is half-max (Fig. 4B,C) – evolves rapidly from *x* ∼ 0.1-0.2 at the start of nc 11 to *x* ∼ *x_c,f_* by the end of nc 13, which occurs around t= 30 minutes (Fig. 4D). Gene expression boundaries then oscillate around *x_c,f_* in both cases through nc 14. To measure which case (Dl-only vs. FFL) results in more stability in nc 14, we once again calculated the ratio *ρ* (see annotation in Fig. 4D). For the parameters chosen in the simulations in Fig. 4B-D (see Methods), we found that the FFL case had less variability than Dl-only case (*ρ* = 0.74 < 1).

To investigate the mechanism by which the FFL may stabilize gene expression boundaries, we analyzed the dynamics of Dl and *Twi* in the FFL case. Within the first 20 minutes of nc 14, Dl concentrations are initially low, then increase rapidly. However, during this time window, *Twi* is not being produced due to the low Dl levels. Thus, as Dl levels are rising, *Twi* levels are falling (highlighted region in Fig 4E), which stabilizes the activation signal (Eqn 9; Fig 4E). This stabilizing effect can be seen by comparing the behavior of the C expression border for the two cases: between ∼35 – ∼45 min, the expression border of C controlled by Dl-only fluctuates much more than the expression border of C controlled by the FFL (highlighted region in Fig. 4D).

To determine whether stabilization by *Twi* can be achieved with a wide range of parameter values, we varied the affinity of Zld for DNA, *K_Zld_*, and the diffusive length scale of Twi, *λ_Twi_*. The lifetime of early zygotic patterning proteins has been experimentally measured to be around 10 minutes (Boettiger and Levine, 2013; Reeves et al., 2012); therefore we set *τ*_Twi_ = 10 min. Theoretically, the higher the cooperativity between Zld and Dl, the more influence Zld has on the Dl system. We found that as the value of cooperativity was increased, generally decreased (see Fig. 4F and Fig. S4). Hence, we chose an intermediate value of *ω* = 30, for further analysis of the model.

When we varied the diffusive length scale of *Twi* (*λ*), setting *K_Zld_* = 1, we found that the ratio *ρ* generally achieves a minimum at intermediate ranges of *λ*, with the exception of targets close to the domain of *Twi* (Fig. 4G). The minimum in *ρ* occurs at higher values of *λ* for gene expression domains further away from the *Twi* boundary and lower values of *λ* for domains closer to the *Twi* boundary. In other words, in order for the *Twi* FFL to affect genes far from the ventral midline, the *Twi* diffusive length scale must be long. However, a longer length scale is detrimental to genes closer to the ventral midline, likely because this spreads *Twi* too thinly. Given this result, and the fact that the *Twi* gradient does not appear to extend to the dorsal midline (Bothma et al., 2018; Zinzen et al., 2006), we expect that *λ* ~ 0.1 is a reasonable estimate of the diffusive strength of Twi.

We then varied the binding affinity of Zld (*K_Zld_*) and found the ratio has a minimum near *K_Zld_* =3 for all genes (Fig. 4H). To determine why *ρ* as a function of *K_Zld_* would have a minimum, we analyzed how the dynamic behavior of *v*, the factor that modulates the effective Dl and *Twi* concentrations (Eqn 1), depends on *K_Zld_*. If the affinity of Zld is too low (*K_Zld_* too high), then the Zld binding is so low that it only weakly modulates the effective Dl concentration (*i*.*e*., *v* ≈ 1 across nc 11-14; green curve in Fig. 4I). Similarly, if the affinity is too high (*K_Zld_* too low), then the DNA is always saturated with Zld, even when Zld reaches its lowest level at the end of nc 14 (*i*.*e*., *v* ≈ *ω* across nc 11-14; blue curve in Fig. 4I). In either case, a constant value of occurs, which implies that the effective concentrations of Dl and Twi, given by *v*[*Dl*] and *v*[*Twi*] respectively, are not appreciably modulated in time, and the beneficial effects of Zld (Fig. 3F) are not realized (see Eqn 10, Methods). On the other hand, if *K_Zld_* is of order 1, the dynamic range of across nc 11-14 is maximized. This is because the normalized concentration of Zld is roughly order 1: it is a constant 1 from nc 10 through the end of nc 13, then decreases to 0.2 by the end of nc 14. In order to counteract the rapid increase in the Dl concentration in nc 14, the effect of Zld must respond by decreasing in magnitude so that the activity remains stable. This occurs when the binding affinity of Zld is of order 1.

### twi-mutant model

In our above analysis, we analyzed two cases: the Dl-only case and the wildtype (FFL) case, in which both Dl and *Twi* regulate C. Since *twi* mutants (Fig. 5A(i)) represent an experimentally attainable model, herein we include a similar analysis comparing the performance of the *twi* mutant to that of the FFL case (see Fig. 5A). It should be noted that the Dl-only case is not a *twi* mutant; it is a hypothetical construct wherein C is designed to only respond to Dl. Thus, the primary difference between the Dl-only case and the *twi* mutant is that in the former, Dl is both necessary and sufficient to independently place gene expression boundary at *x_c,f_*, whereas in the latter, the loss of *Twi* results in a ventral shift in gene expression boundary.

**Figure 5.**
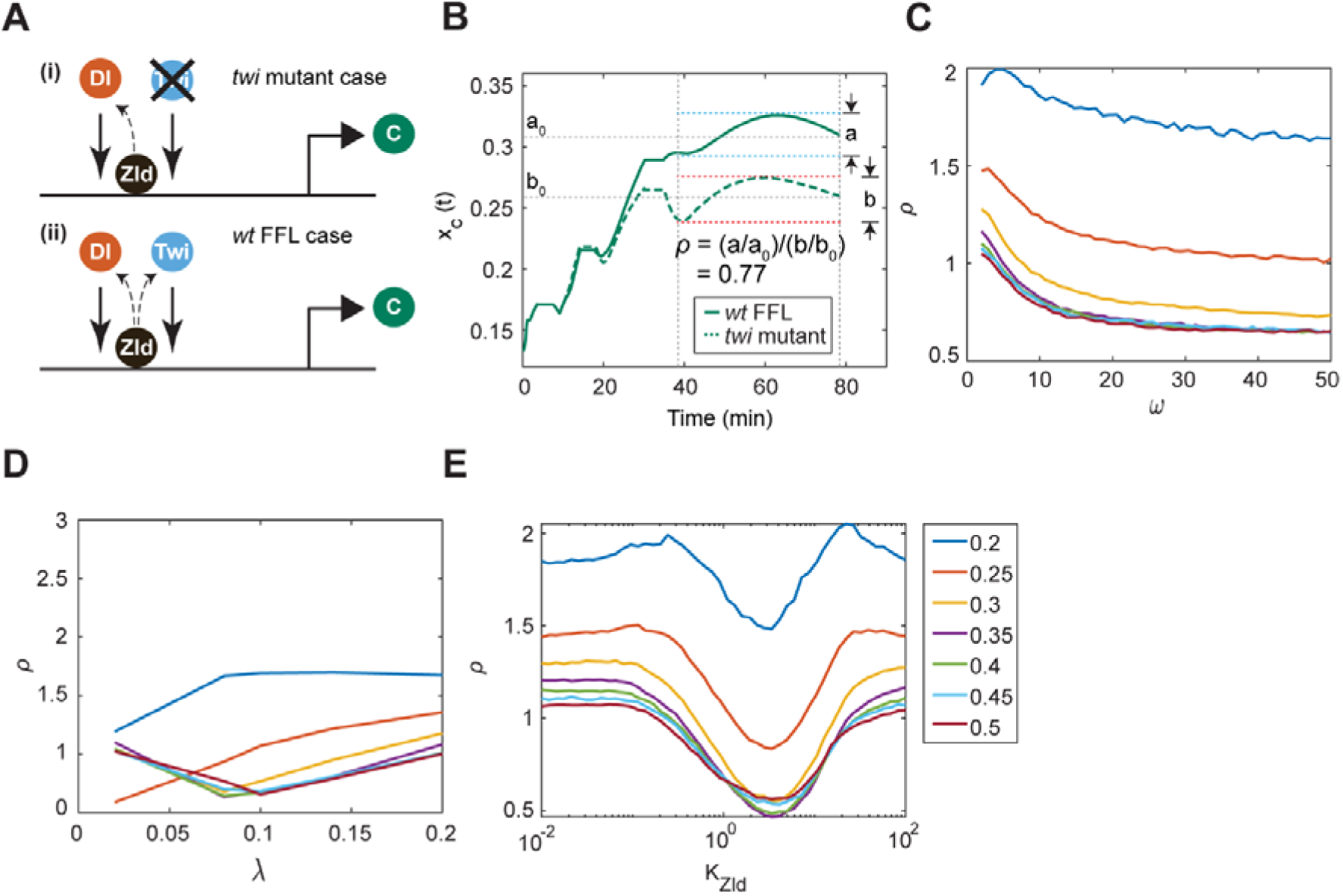
Effect of Dl-Twi FFL in the *Twi* mutant model. **A.** The enhancer module of C in the case of (i). *twi* mutant and (ii) wildtype FFL. Both cases include the effect of Zld. **B.** Variation in the boundary of C against time for both cases. The extent of variation is normalized by the final value of x_C_ (t). **C.** Variation of the extent of oscillations with at different values of *x_c,f_* **D.** Variation of the extent of oscillations with at different values of *x_c,f_* **E.** Variation of the extent of oscillations with *K_Zld_* at different values of *x_c,f_*

In our analysis, we found that in many cases, wildtype gene expression boundaries were more stable than those in the *twi* mutant (Fig. 5B). It should be noted that, since the final expression boundaries for the wildtype vs. *twi* mutant cases are different, we slightly modified our metric *ρ* (see Methods for details).

To determine whether our observation – that the FFL stabilizes gene expression with respect to the *twi* mutant case – is general, we varied the *Twi* diffusive length scale, the affinity of Zld, and the cooperative strength of Zld. Similar to our comparison of Dl-alone vs the wildtype FFL, we found that with increasing cooperativity between Zld and Dl, the value of *ρ* generally decreased for all values of *x_c,f_*. Therefore, we set an intermediate value of *ω* = 30 for further analysis (see Fig. 5C and Fig. S5). We also found that genes further away from the ventral midline are benefitted by longer diffusive length scales, and vice versa (Fig 5D). However, the curves were generally shifted upwards (compare curves in Fig. 5D to those in Fig. 4G), which perhaps surprisingly implies that the *twi* mutant is slightly more stable than the Dl-alone case. We also found similar behavior of *ρ* as a function of *K_Zld_* (Fig. 5E; cf Fig. 4H) and of *ω* (Fig. 5C; cf Fig. 4F), each with the added upward shift. In general, regulation by Dl/Twi FFL appears to be more stable than regulation by Dl in *twi* mutants, which in turn is more stable than regulation by Dl when it acts independently. The exceptions are genes close to the *twi* expression boundary, which are almost always more stable in *twi* mutants than in wildtype, unless the *Twi* diffusion length is very short (Fig. 5D). Besides those exceptions, the general behavior may be attributed to the ventral shift of gene expression in *twi* mutants, which places gene expression in regions of steeper Dl gradient slope. A steeper slope implies that variation in Dl concentration can be absorbed better, in that the location where Dl achieves a given threshold has less variation (Reeves et al., 2012). This effect is prominent closer to the ventral midline and wanes away from it, as binding affinities become similar.

## Discussion

Feedforward loops are one of the most commonly-found – and extensively-studied – motifs in a wide variety of biological networks, such as transcription factor networks, metabolic pathway networks, gene regulatory networks, signaling networks etc. (Davidson et al., 2002; Duggan et al., 1998; Mangan et al., 2003; Schmitz et al., 2011; Shen-Orr et al., 2002; Sunaga et al., 2019). Most quantitative, theoretical, or computational studies of FFL function have focused on purely time-varying systems, such as single-cell organisms, although a handful of studies have introduced a spatial component (Hironaka and Morishita, 2012; Schaerli et al., 2014). In this work, we have analyzed the function of a coherent FFL in a non-steady-state, spatially-distributed morphogen system. First, we used a model with a generic morphogen to show the versatility of an FFL. Next, we used spatiotemporal data of Dl in the early *Drosophila* embryo as the primary morphogen in a realistic model to extend the analysis of FFLs to a real biological network. In both cases, we find that there are viable regions in the parameter space of both models that make an FFL network motif a robust choice for development.

In the generic morphogen gradient model, we found that the effectiveness of the FFL was largely independent of the parameters of the model. From previous analyses of coherent FFLs, primarily in well-mixed systems, we would expect that small fluctuations in the gradient of the primary morphogen would be buffered by the intermediate node (Mangan and Alon, 2003). A wide range of time and length scales were explored in the model and in almost all cases, regulation by an FFL resulted in robust expression boundaries, in the sense of lesser oscillations over time, throughout the tissue domain. We found that two aspects in particular contributed to the increased stability in the output gene. Most importantly, the lifetime of the intermediate transcription factor was required to be roughly as long as (or longer than) the fluctuations in the input transcription factor. This fact allowed the intermediate to smooth out the fluctuations in the input, roughly acting as a noise-filtering mechanism. Second, the output gene could be more stabilized if the fluctuations in the intermediate were shifted slightly out of phase with the input fluctuations. In this case, the oscillations in the intermediate would somewhat cancel out the oscillations in the input. Given that these two aspects arise in our system under a wide range of parameter values, the superior performance of the spatiotemporal FFL can be attributed to the structure of the network motif rather than a result of any particular choice of model parameters.

We extended this model to use the dynamic profile of Venus-tagged Dl as the primary morphogen and included the dynamics of the target gene in a more realistic model (Reeves et al., 2012). The major oscillations in the Dl gradient are due to nuclear divisions, and, unlike in the generic model, these divisions are limited in time (only four nuclear divisions occur in the relevant stage) and do not have a regular, repeated period. The last nuclear cycle is about four times longer than the previous nuclear cycle and a substantial increase in the concentration of Dl is observed. The growing amplitude, limited number of oscillations, and short duration of nc 10-13 interphases implied that, prior to nc 14, gene expression may be limited. Indeed, when we used the Dl/Twi FFL to place the domains of target genes, we observed that expression levels of all Dl targets, including that of the intermediate node of the FFL, Twi, were insignificant until nc 14. This problem had been observed in previous models that tried to simulate gene expression domains of Dl targets and could not explain the lack of expression in the previous nuclear cycles (O’Connell and Reeves, 2015; Reeves et al., 2012). However, gene expression for many Dl targets has been experimentally observed as early as nc 12 (Liang et al., 2008; Nien et al., 2011; Ozdemir et al., 2011; Sandler and Stathopoulos, 2016; Sandmann et al., 2007). The ubiquitous pioneer factor Zld has been shown to coordinate the temporal activation of transcription by binding to DNA as early as nc 2 (Nien et al., 2011). In Zld mutant embryos, several genes either fail to activate or are delayed, with some showing sporadic expression. In addition, Zld binding sites have been found in large numbers near transcription start sites of several of these genes as well as near their enhancer modules (Nien et al., 2011; Sun et al., 2015). Zld has also been known to increase the local concentration of patterning transcription factors such as Bcd and Dl, thus effectively increasing its local activity (Dufourt et al., 2018; Mir et al., 2017; Yamada et al., 2019). Considering these known effects of Zld, we decided that Zld was a crucial and well-founded missing piece of our model.

The role of Zld in early transcriptional activation was seen due to an increase in the activity of Dl in earlier nuclear cycles, allowing *twi* to be expressed in late nc 12 (see Fig. 3D,F). Furthermore, the role of Zld in stabilizing target gene expression occurs due to opposite trends in Zld and Dl concentrations in nc 14. The rapid increase in Dl concentrations in nc 14 was counteracted by a decrease in the concentration of Zld, resulting in a relatively stable activation signal, which translated to stable gene expression (see Fig. 3D,F,). These two effects, coupled together, resulted in early gene expression that was more stable than the case when effects of Zld were not included.

As with the generic model, the stability of gene expression borders can be at least partially attributed to the phase difference between Dl and *Twi* oscillations: at the onset of nc 14, Dl levels are increasing while *Twi* levels are decreasing. This results from the fact that, at the boundary between nc 13 and 14, the Dl gradient sharply decreases in amplitude, which momentarily halts *Twi* production. Thus, the FFL activation signal is only partly disturbed due to the mitosis event between nc 13 and 14, due to lag in the response of Twi. This phase difference fades later in nc 14 when both Dl levels continue to increase, and production of *Twi* resumes. The degree to which *Twi* levels declined at the onset of nc 14 was dependent on the specific values of the model parameters, but the duration was largely independent of the parameters chosen. Therefore, this phase difference, and thus the robustness of the FFL-dependent gene expression, is mainly determined by the wiring of the network motif itself.

Given the stability of gene expression that is imparted to the system by the FFL, one would expect *twi* mutants to be less stable than wildtype. Indeed, literature shows that the border of *sna* is less well-defined in *twi* mutant embryos, which could imply a less stable *sna* border (Ip et al., 1992b). While the model does in fact predict that some DV gene expression in *twi* mutants is less stable, the increased sensitivity is not as great as one might expect. In fact, the *twi* mutant case outperformed the Dl-only case. This is possibly due to the fact that gene expression is shifted ventrally in *twi* mutants (*i*.*e*., a gene with a border located at 0.3 in the wildtype FFL case may shift to 0.25 in the *twi* mutant). More ventral positions experience a steeper Dl gradient slope, which in turn intrinsically imparts more robustness to gene border locations with respect to fluctuations in morphogen concentration (Reeves et al., 2012). The end result is that some of the increase in sensitivity due to loss of *Twi* activity is partially counterbalanced by the ventral shift in the gene of interest. Even so, the model predicts that, for reasonable ranges of parameter values, *twi* mutants would indeed have more dynamic gene expression borders. This prediction could be tested by quantitative imaging of Dl-target gene dynamics in live, *twi* mutant embryos.

Coherent FFLs have been shown to buffer abrupt changes in the input signal (Mangan and Alon, 2003). We observed this phenomenon under general conditions in both the generic, oscillating morphogen model and in our Dl/Twi/Zld model, where abrupt changes in the morphogen concentration of Dl, due to mitosis events, are buffered by Twi, ensuring that the activation signal perceived by the target genes is stable. We predict that other dynamic, morphogen-mediated systems may have cFFLs that stabilize gene expression borders.

## Methods

### Generic morphogen model details

In this model the primary morphogen A, was treated as a generic transcription factor, whose protein concentration oscillates sinusoidally in time and decays exponentially along a single spatial dimension which is varied from 0 to 1:

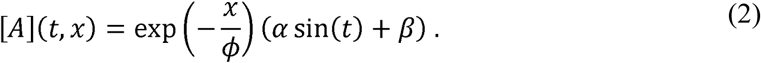

In our model, we chose *ϕ* = 0.2, *α* = 0.1, and *β* = 1. The primary morphogen activates the secondary transcription factor B at a threshold that, on average, corresponds to a gene expression border at *x* = 0.2. The dynamics of the intermediate node B is described by the following differential equation,

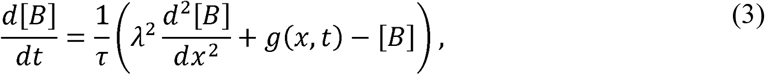

where [B] refers to protein concentration of B, *λ* represents diffusion constant of B, *τ* represents lifetime of B and *g*(*x,t*) is a smoothened step activation function which activates expression for a threshold in A that corresponds, on average, to *x* < 0.2. Either A alone, or both A and B together, activate(s) the output target gene (C).The gene expression border of C, *x*_*c*0_, was defined as the location of the border when a= 0, and was chosen to vary between *x* = 0.2 and *x* = 0.5. The location of B was chosen to roughly coincide with location of the boundary of Twist in the DV axis of the early *Drosophila* embryo. We evaluated the stability of the gene expression border of C in two cases – 1). When protein A independently controls the expression border of gene C (“A-only”), and 2). when the gene expression border of C is jointly regulated by A and B using OR-gate logic in a feed forward loop (“FFL”).

The concentration profile of target C, in the A-only case is described by the following Hill function,

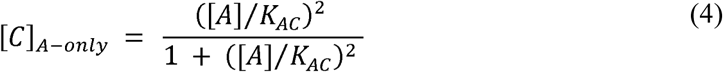

and in the FFL case is described using an FFL with OR-gate logic as follows,

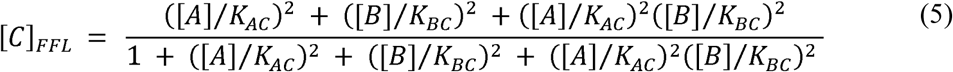

where *K_ij_* represents binding affinity of factor *i* to the enhancer regions of *j*. The affinities *K_AC_* and *K_C_* were set equal to each other and were tuned so that *x*_*c*0_ would take on a chosen value between *x* = 0.2 and *x* = 0.5 at *t* = 100 min.

### Dl/Twi/Zld model details

In this model, we replaced the generic morphogen with experimentally derived expression data from Venus-tagged Dl (Reeves et al., 2012). The simulation was run for the length of the time period between nc 10 and nc 14, which was roughly 80 minutes. The spatial dimension is scaled to vary between 0 and 1, extending from the ventral to the dorsal midline. As with the generic morphogen model, Dl activates *Twi* and several other targets, referred to herein as C.

We evaluated the stability of the gene expression border of C in two cases – 1). When Dl protein independently controls the expression border of C (“Dl-only”), and 2). when the gene expression border of C is jointly regulated by Dl and *Twi* using OR-gate logic in a feed forward loop (“wildtype FFL”).

The dynamics of *Twi* is described by the following differential equation,

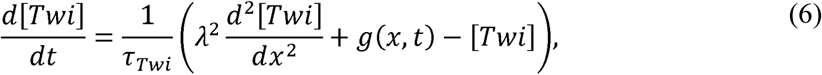

where [Twi] refers to a lumped variable of the mRNA or protein concentration of Twi, *λ* represents diffusion constant of Twi, *τ* represents lifetime of Twi, and *g*(*x,t*) is a smoothened step activation function which activates *Twi* at a threshold of Dl that corresponds to roughly *x* < 0.2 at the end of the simulation (end of nc 14). Dynamics of C is incorporated in this model, which is given by the following differential equation,

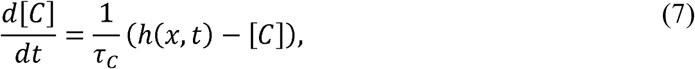

where [C] refers to mRNA concentration of C, *τ*_C_ refers to the lifetime of C and h(x,t) is the activation function given by the following Hill function,

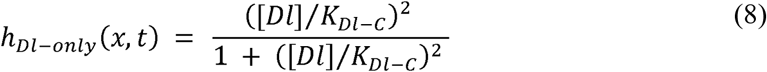

when the dynamics are controlled independently by Dl, and by the following Hill function

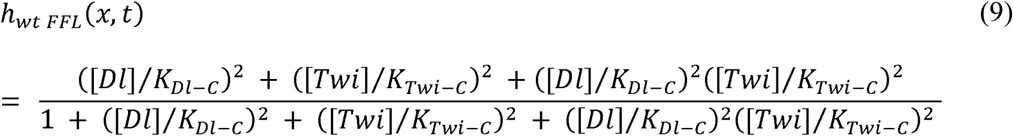

when dynamics are controlled by a Dl-Twi FFL governed by OR gate logic. The affinities, *K _l-C_* and *K_Twi-C_*, were set to be equal. The affinities were tuned so that the expression boundary for *C, x_C_* (*t*), would take on a chosen value, *x_c,f_*, between *x* = 0.2 and *x* = 0.5 if the final Dl gradient and Zld concentration at the end of nc 14 were held constant and both *Twi* and C came to steady state.. Note that, by definition, the value of binding affinity *K_Dl-C_* would be different in the two cases.

We included the effect of ubiquitous factor Zld that has the following variation in time,

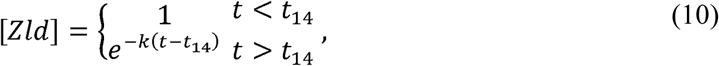

where, is *k* a constant and *t*_14_ is the time of the beginning of nc 14. The value of *k* was chosen so that zldl reached 0.2 by the end of nc 14. Zld influences the activity of Dl by changing its effective concentration to nll, where is as defined in Eqn 1.

### twi mutant case

The *twi* mutant case had the same formulation as the Dl-only case; however, the binding affinity, *K_l-C_*, in the *twi* mutant model was set to be equal to that in the wildtype FFL case. In the FFL case, the binding affinity was tuned, together with *K_Twi-C_*, to ensure the gene expression boundary was placed at *x_c,f_* (see above). In the *twi* mutant case, *Twi* activity is lost, but the Dl binding affinity remains the same, which ensures that gene expression in the *twi* mutant would be shifted ventrally as compared to the wildtype FFL case (*i*.*e*., *x_c,f_* is different between the two cases). This is in contrast to the comparison between the Dl-only case and the wildtype FFL case, in which the binding affinities are intentionally different between the two cases so that *x_c,f_* is held the same.

In addition, since the final gene expression boundary location in the *twi* mutant case is different from the FFL case, we use the following metric to compare stability of oscillations in the boundary

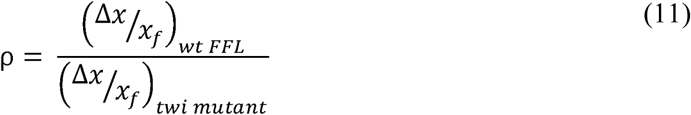

such that, 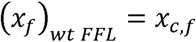

