## Supplemental Methods and Figures for "Spatiotemporal control of gene expression boundaries using a feedforward loop"

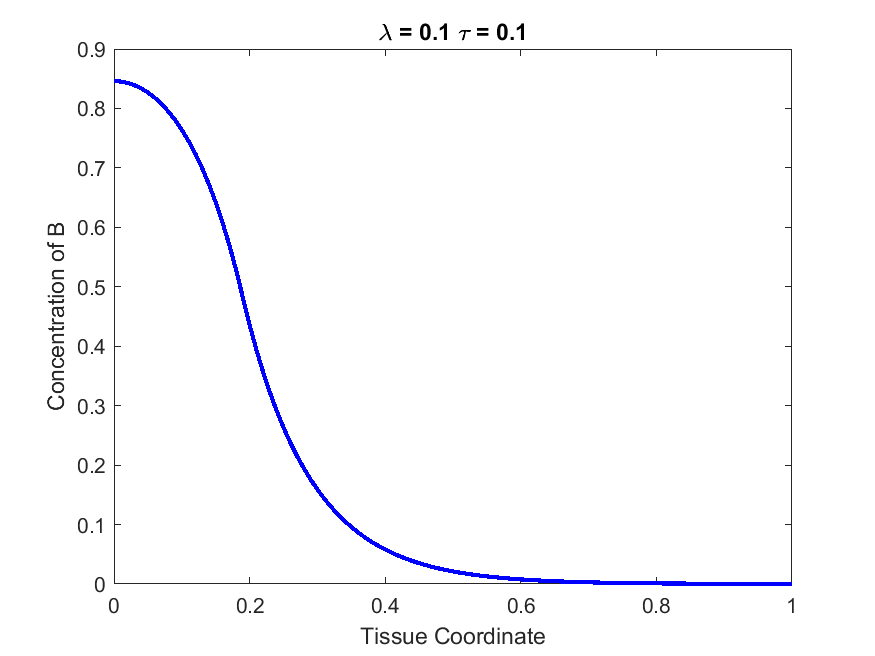

Figure S1. Concentration of B varying with Tissue Coordinate at the final time point for parameters $\lambda= 0.1$ and $\tau= 10$.

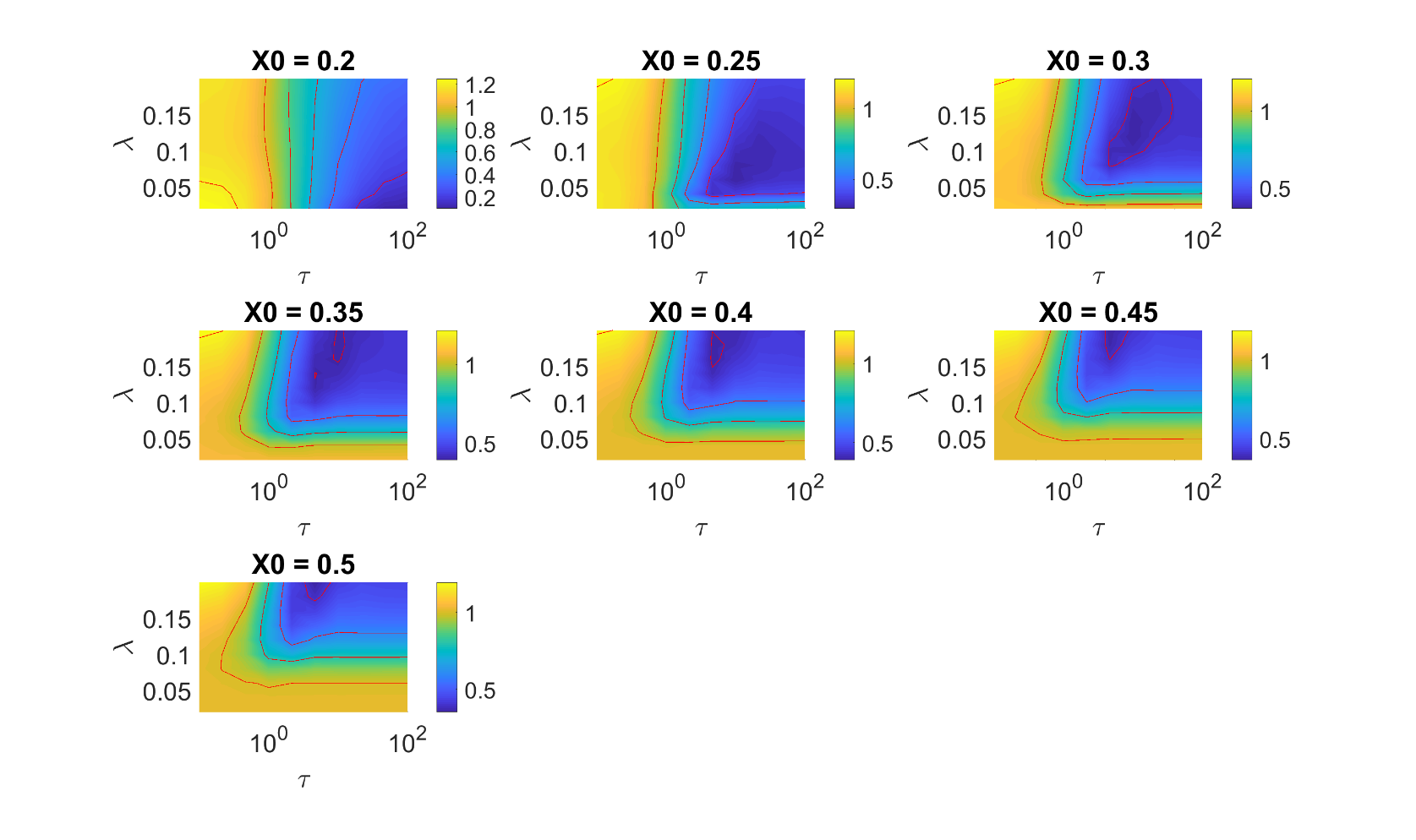

Figure S2. Contour plots of $\rho$ as a function of $\lambda$ and $\tau$ for the generic model.

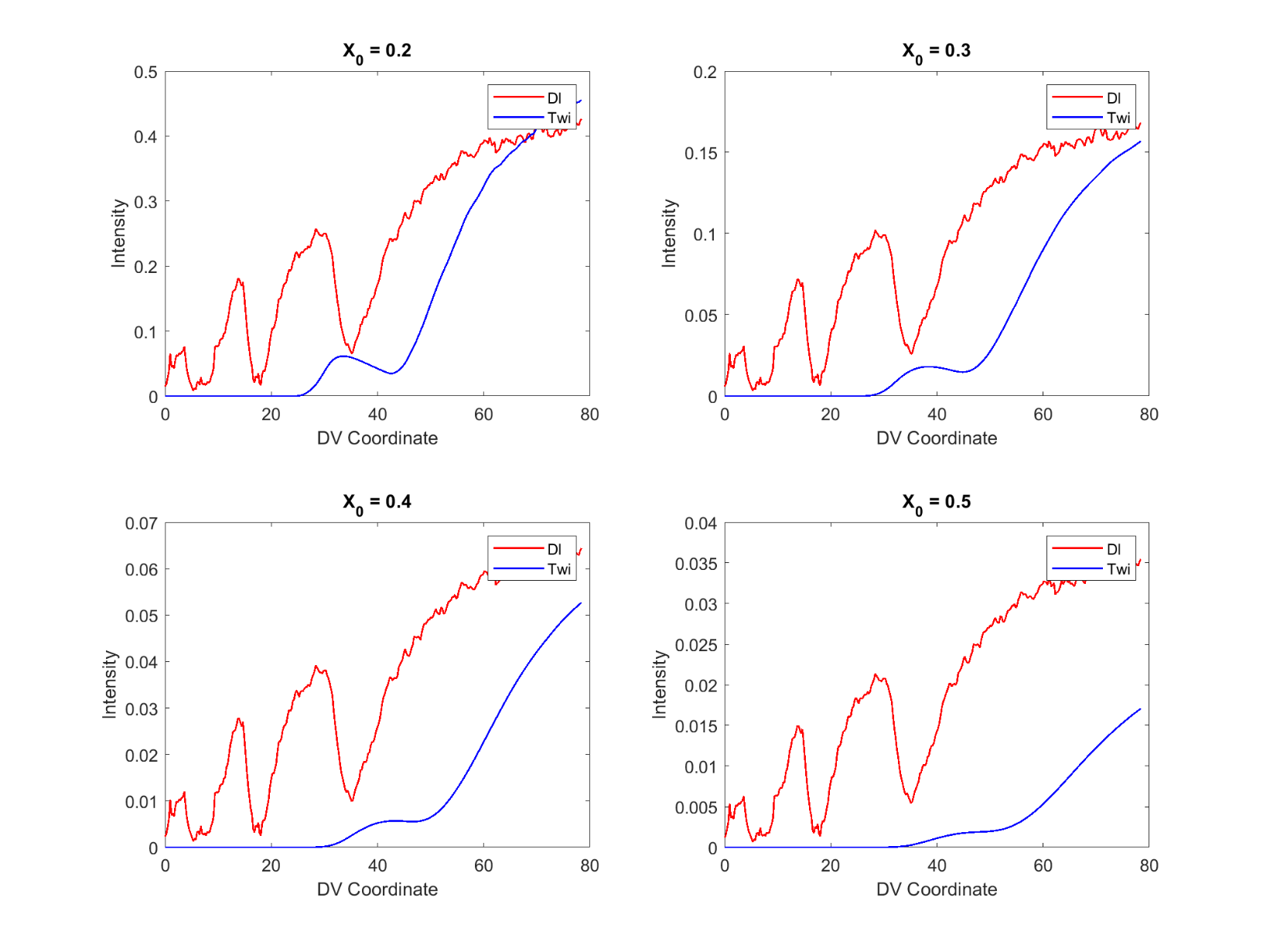
Figure S3. Dl and Twi concentration curves at different values of the DV coordinate. ($\lambda= 0.1,$ $\tau= 10$)

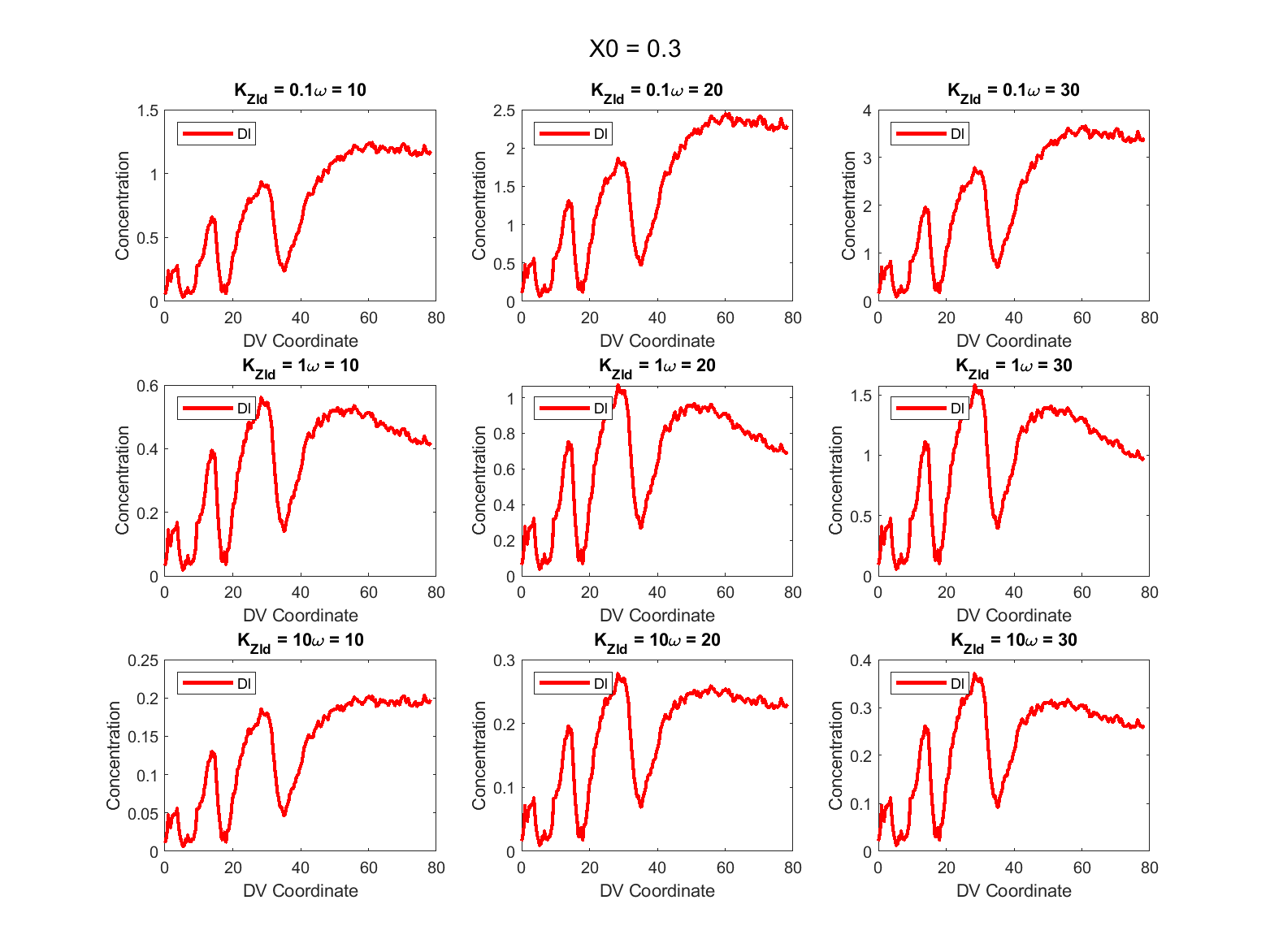

Figure S4. Effect of Zld on Dl for varying parameters of $K_{Zld}$ and $\omega$ at $x_{0}=0.3$, $\lambda= 0.1$ and $\tau= 10$.

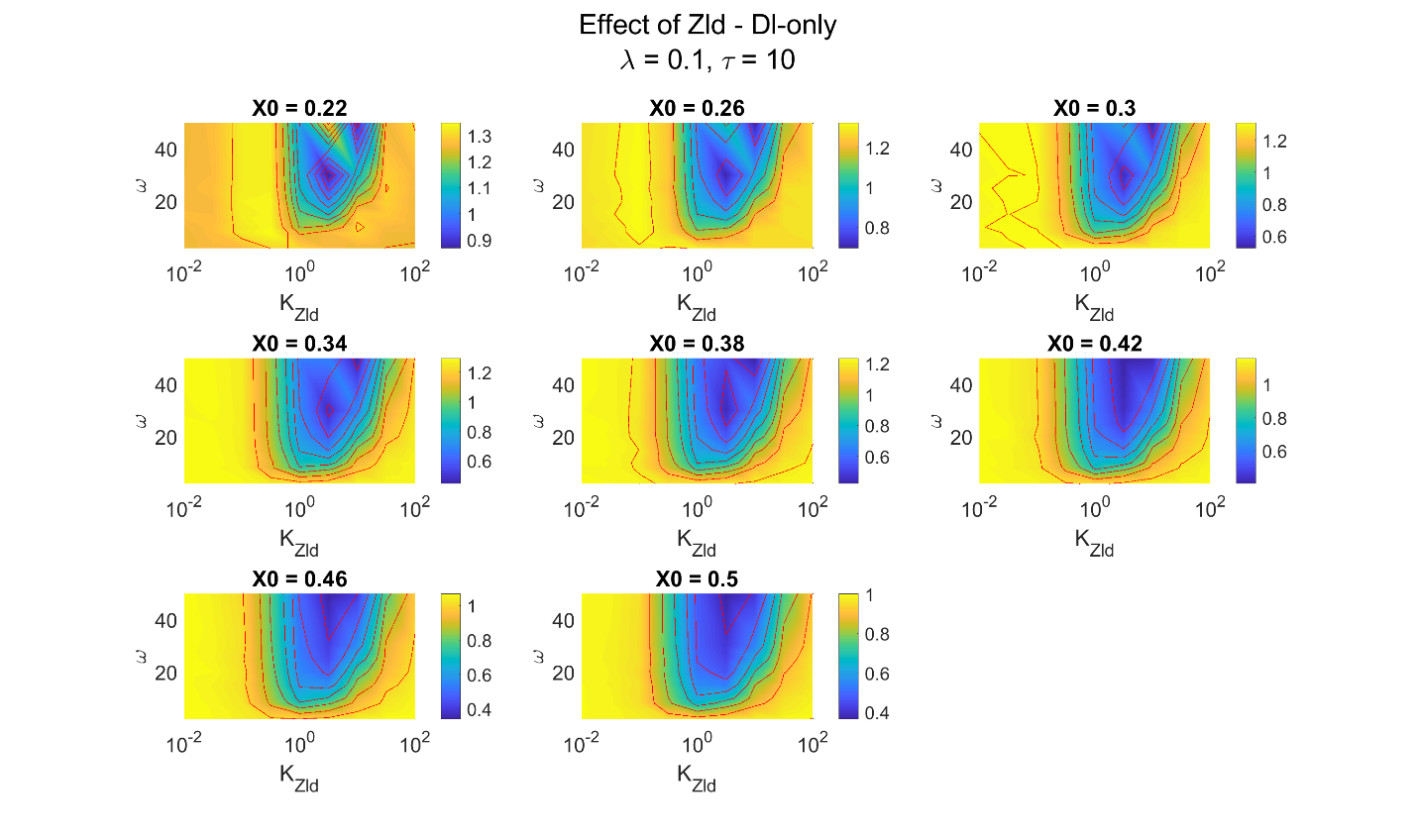
Figure S5. Contour plots $\rho$ as a function of $K_{Zld}$ and $\omega$ for the Dl/Twi/Zld model for different values of the boundary comparing *wt* FFL and Dl-only.

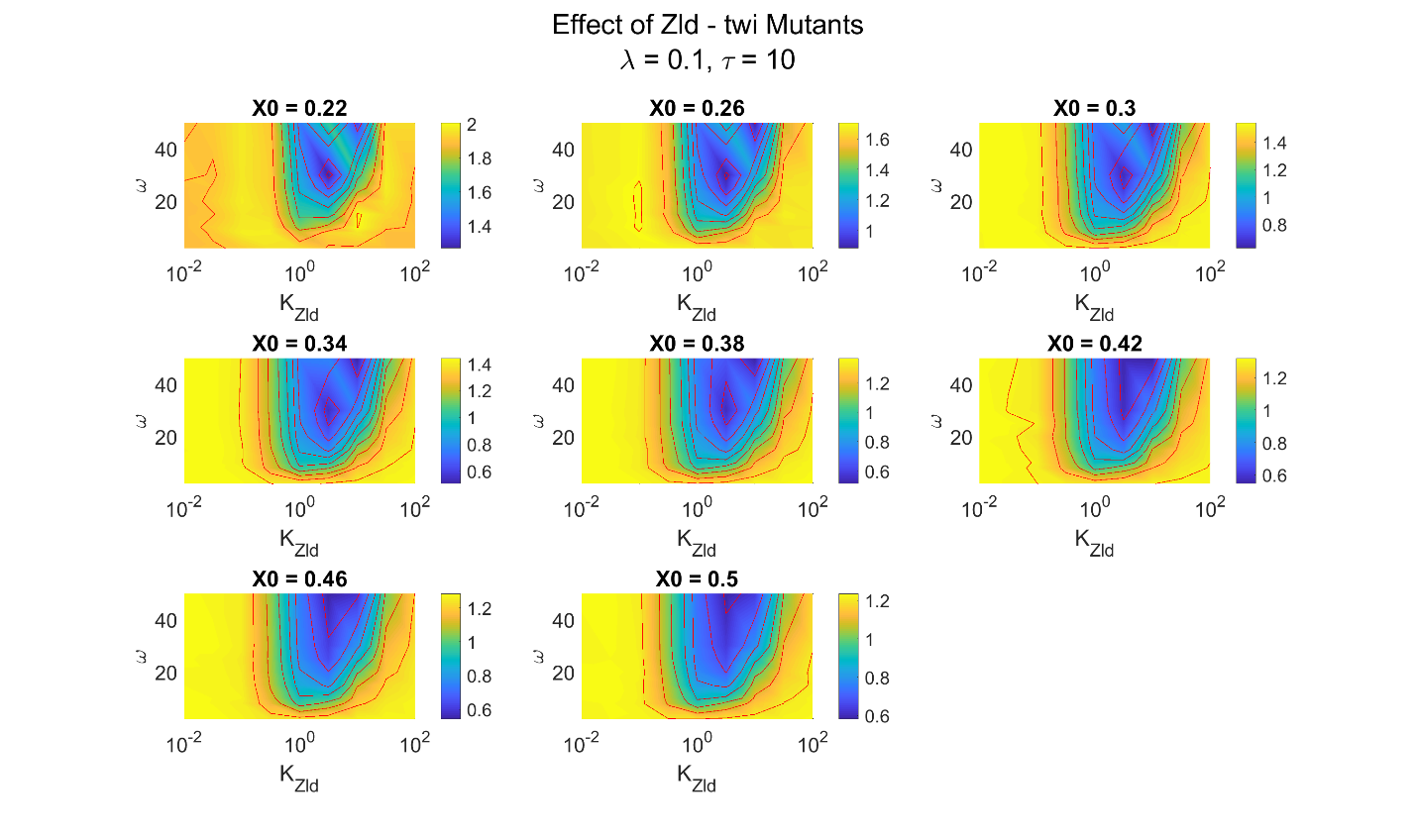

Figure S6. Contour plots $\rho$ as a function of $K_{Zld}$ and $\omega$ for the Dl/Twi model for different values of the boundary comparing *wt* FFL and *twi* mutants

**Supplemental Movie Legends.**

**Movie 1**. The movie shows the variation in concentration of A (in red), B (normalized; in blue) and, C (normalized) in the two cases – “A only” (in green; dashed curve) and “FFL” (in green; solid curve) at $\lambda= 0.1$ and $\tau= 0.1$. In this movie, the all concentration curves seem to oscillate in phase with the oscillations of A.

**Movie 2.** The movie shows the variation in concentration of A (in red), B (normalized; in blue) and, C (normalized) in the two cases – “A only” (in green, dashed curve) and “FFL” (in green; solid curve) at $\lambda= 0.1$ and $\tau= 1$. In this movie, the higher lifetime of B contributes not only to its own stability but also to the stability of C in the FFL case. Note the slight phase difference in the oscillations of A and B which is clearly observable near the boundary of B ($x = 0.2$) and is less clear elsewhere.

**Movie 3.** The movie shows the variation in concentration of A (in red), B (normalized; in blue) and, C (normalized) in the two cases – “A only” (in green; dashed curve) and “FFL” (in green; solid curve) at $\lambda= 0.1$ and $\tau= 100$. In this movie, the lifetime of B is very high, and it oscillates minimally. This has a noise filtering effect on C and its concentration profile can be seen oscillate lesser in the FFL case.
